# Periodicity, Mixed-Mode Oscillations, and Multiple Timescale in a Phosphoinositide-Rho GTPase Network

**DOI:** 10.1101/2023.05.22.541800

**Authors:** Chee San Tong, Min Wu

## Abstract

While rhythmic contractile behavior is prevalent on the cortex of living cells, current experimental observation and mechanistic understanding primarily tackle a small subset of dynamical behavior including excitable or periodic events that can be described by simple activator-delayed inhibitor mechanisms. In this work we found that the oscillatory activation of Rho GTPase in nocodazole-treated mitotic rat basophilic leukemia (RBL) cells exhibited both simple and complex mixed-mode oscillations, with periodicity ranging from 30 sec to 5 min. Complex mixed-mode oscillations require at least two instability-generating mechanisms. We show that Rho oscillations at the fast timescale (20-30 sec) is regulated by phosphatidylinositol (3,4,5)-trisphosphate (PIP3) via an activator-delayed inhibitor mechanism, while the period of the slow reaction (minutes) is regulated by phosphatidylinositol 4-phosphate (PI(4)P) via an activator-substrate depletion mechanism where replenishment of phosphatidylinositol (4,5)-bisphosphate (PI(4,5)P2) is rate-limiting. Conversion from simple to complex oscillations could be induced by modulating PIP3 metabolism or membrane contact site dynamics. In particular, a period-doubling intermediate can be captured by PTEN depletion. Both period doubling and mixed-mode oscillations are intermediate states towards chaos. Collectively, these results suggest that phosphoinositide-Rho GTPase signaling network is poised at the edge of chaos and small parameter changes in the phosphoinositide metabolism network could confer cells the flexibility to rapidly transit into a number of ordered states with different periodicities.

## Introduction

Spatiotemporal regulation of cell contractility is important for most morphogenesis events including cell division. Contractile forces are generated by actomyosin networks through the activity of myosin II motors that slide antiparallel actin filaments relative to each other. Assembly of actomyosin network is in turn governed by Rho GTPase activity. In mitosis, Rho is necessary for recruitment and assembly of cytokinetic ring components in animal cells, followed by ring contraction and cell division. How Rho is recruited to and regulated on the plasma membrane of mitotic cell is not completely understood (1). In addition, nonlinear dynamical behaviors associated with Rho dynamics, such as waves, pulses, oscillations, or wandering furrowing activity, has been reported for various mitotic cell types, including frog and echinoderm oocytes and embryos (2-5), C. elegans embryos (6-8), Dictystelia (9), RBL mast cells (10), and epithelial cells (11, 12). These observations increasingly support the idea that the contractile machinery for cell division is composed of local furrowing units that can independently assemble, disassemble, pulse, or oscillate, and that the coordination of these contractile units lead to formation of productive furrowing, as opposed to a single coherent contractile band that function as a ‘purse string’ (13). Understanding the onset, frequency, duration, amplitude and coordination of these contractile units and the underlying nonlinear network structure governing such dynamics will thus be fundamental to our understanding of cytokinesis.

For practical reasons, mechanistic understanding of Rho signaling network topology has mostly come from studies of interphase cells where surface contraction pulses or oscillations are also prevalent. The classic model describes contractile pulses by focusing on the mechanical property of the actomyosin network, or the “cytogel”, in a manner analogous to how Hodgekin Huxley equations describe action potential with a single measurement of membrane potential (14). These models were inspired by works on model systems of large sizes such as the true slime mold Physarum polycephalum where cycles of cytoskeletal expansion and contraction can be readily observed and mechanical properties can be measured (15). Actomyosin-centric mechanical models were also supported by recent experimental work (16-18). However, information on the chemical state of the “cytogel”, also essential ingredients in the mechanochemical model, is largely absent. Probing the chemical state of the system only became possible in recent years with fluorescent reporters to directly visualize Rho and its interaction networks (4, 19, 20). With quantitative investigation of the behavior of RhoA together with actomyosin networks in the contractile pulses and experimental perturbation to these patterns, many studies suggest, perhaps surprisingly, that oscillatory cycles of Rho persisted in the absence of myosin II (8, 21). An emerging view is that there should be an excitable Rho GTPase network corresponding to cycles of Rho GTPase activity that acts upstream of myosin II and drive cycles of contraction (22). Nevertheless, dissecting such network experimentally remain challenging.

The generic principle of oscillations requires the interplay of a local self-enhancement activator and an antagonistic effect that decreases production of activator or increases its degradation. A number of studies have proposed that RhoGEF and RhoGAP could play the role as the activator and inhibitor (2, 7, 19). Some open questions remain. First, the identity of specific GEFs or GAPs differ in different systems (22). This may seem natural because mapping the topology of signaling networks corresponds to solving an inverse problem and inverse problem tend to have multiple solutions. Complications also exist with regard to what is sufficient to cause changes in Rho pulses does not appear to be absolutely necessary (23). Second, mechanisms regulating the Rho oscillation period, or refractory phase, are not well understood. Rho pulses in most systems are highly irregular and heterogenous. These experimental data likely came from the excitable regimes that do not give enough constraints to falsify models because interpeak interval here is not determined by the duration of refractory phase. In experimental conditions that generate robust oscillation, oscillation periods are highly variable both within a single system and between systems. Reported values include 80 – 120 sec in starfish oocytes and activated frog eggs (4), 50-120 sec (5), and 30 – 50 sec in U2OS cells (19). In addition, multiple harmonics have been reported in Physarum polycephalum (24). The most compelling evidence where perturbation of a localized inhibitor could modulate Rho oscillation period came from study of Xenopus oocyte where dose-dependent increase of RhoGAP RGA-3/4 leads to a corresponding decrease in oscillation period from 150-250 sec to 50-100 sec (23). Currently how RGA-3/4 is recruited to membrane or how duration of RGA-3/4 is controlled is not known. Many of the components in the RhoGEF/GAP network including Rho itself can be regulated by phosphoinositide (25-28). Oscillations of various phosphoinositides have been observed in many single cell systems (29-37), but whether phosphoinositide oscillations are coupled with Rho oscillations, or how phosphoinositides specifically affect Rho signaling networks in general, is not known.

In this paper, we show that Rho oscillations in nocodazole-treated mitotic RBL cells exhibit periodicity ranging from 30 sec to 5 min, which covers periodicities for contractile pulses reported in all experimental systems. We found that the heterogeneity in oscillation frequency is not achieved by a single instability-generating mechanism but requires at least two layers of controls operating at different timescales. We further demonstrate that phosphoinositide metabolism critically controls the timescales for each layer and coupling of two feedback loops generates complex oscillation dynamics such period doubling and mixed-mode oscillations. Our findings expand the rich spectrum of dynamical behavior generated by phosphoinositide-Rho GTPase network and provide an experimental framework to understand heterogeneity in contractile instability.

## Results

To understand the molecular network that controls cell contractility, we looked for experimental conditions that allowed us to generate and perturb Rho oscillations. Oscillators convey systems-level characteristics that is informative for dissecting the full topology of the reaction network (38). For activity sensor of Rho, we use the rhotekin G-protein binding domain (rGBD) which specifically binds to active Rho (RhoGTP) and translocates to the plasma membrane after binding, which can be monitored as increases in fluorescence intensity as visualized by the total internal reflection microscopy (TIRFM). In RBL cells, we found oscillation of Rho dynamics was the most robust when nocodazole, a microtubule depolymerization drug was introduced to mitotic cells (n=29 experiments, 57/70 cells). The effect of microtubule poison on contractility was known in the literature, likely indicating that microtubule has a global inhibitory effect on contractility (39, 40). Rho oscillations in mitotic cells are heterogenous and showed variations both in their periodicities and pulse durations (**Fig 1 A-D**). Based on fast Fourier transform (FFT), the major period of these oscillations varied from ∼ 30 sec to 5 min. The fastest Rho oscillations around 30 sec were rare (8/57 cells) (**Fig 1A**). Most Rho oscillations have lower periodicities (70-180 sec) (30/57 cells) (**Fig 1B-C**). Despite the differences in periodicities, the profile of individual Rho pulses has very similar rise phases. When we aligned the peaks of individual oscillatory cycles with different frequencies it is evident that the rate of rho recruitment in all three types of simple Rho oscillations are similar and can be superimposed. The fastest oscillation has a symmetrical rise (∼16 – 20 s) and fall phases (∼16 -20s) (**Fig 1A**), while Rho oscillations with period of ∼120 sec have similar rise phases but slower decay phase (∼36-44 sec) (**Fig 1B**). Duration of the peak do not linearly correlate with oscillation period, as oscillation with similar period of ∼150 sec can have very different duration as seen from both autocorrelation analysis and the rise/decay phase of averaged profiles (compare **Fig 1B and 1C**).

**Fig 1.**
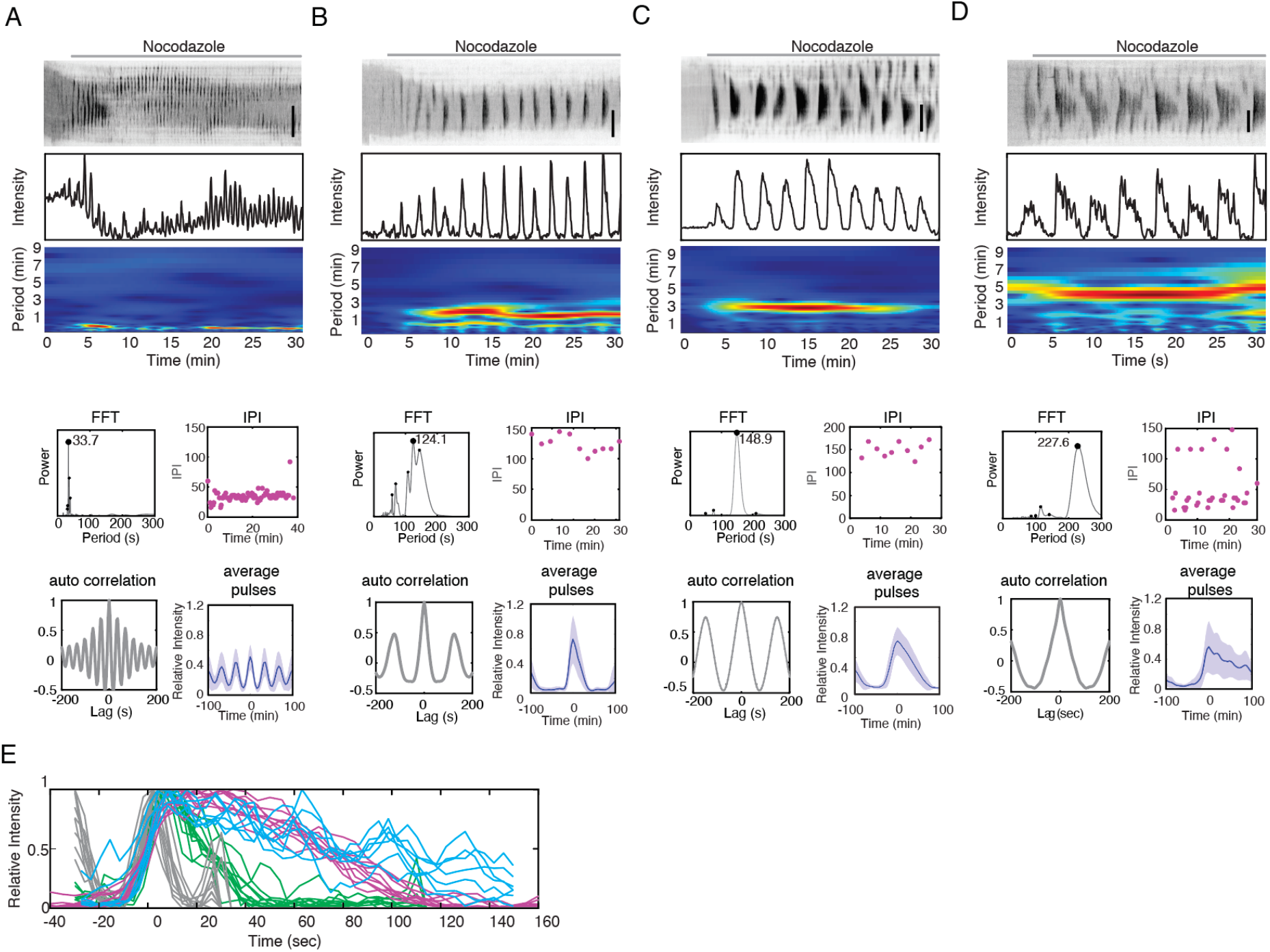
Simple and complex Rho oscillations are observed in mitotic RBL cells treated with nocodazole. (**A-D**) Top: Kymographs, intensity profiles, and wavelet analyses of representative movies with simple and complex Rho oscillation in WT cells, as observed by Rho sensor (rGBD) (n=29 experiments, 57 cells). Bottom: Analyses of properties of Rho oscillations. FFT analysis shows the major period. Interpeak intervals of the representative movies were plotted with respect to changes to time. A, B and C show a single interpeak interval. D shows 2 distinct interpeak intervals. Auto-correlation of Rho oscillations signals shows the duration of peaks. Average profiles by aligned peaks shows the rise and drop phase of an average pulse’s intensity profiles. The solid lines represent the mean intensities of the profile and shaded region represents the standard deviations of the intensities. (**E**) Alignment of individual peak traces of Rho oscillations. (A)=gray, (B)=green, (C)=magenta, (D)=cyan. Vertical scale bar: 10 μm.

The heterogenous oscillation periods and pulse durations make it challenging to identify the negative regulators responsible for controlling these parameters because any conclusions would require large sample sizes for statistical confidence. We therefore took an alternative approach. We noticed that besides simple oscillations, complex mixed-mode Rho oscillations were occasionally observed (**Fig 1D**). Mixed-mode oscillations have oscillatory cycles that consist of large-amplitude oscillations intercalated with small-amplitude oscillations (41). While FFT of these complex oscillations also give rise to a single major period around 5 min, interpeak interval (IPI) analysis reveals two population of the intervals, centered around approximately 120 s for larger gaps and 30 seconds for the ‘nested’ peaks (**Fig 1D**). Complex oscillations arise when oscillators are chemically coupled in tandem, such as when a flow system containing two allosteric enzymes for which the product of one serves as the substrate of the other (42, 43), or in parallel, such as two enzymes catalyzes the same metabolic reaction but with different kinetic (44). These coupled systems exhibit a far richer range of dynamical behavior than either of the two uncoupled systems and offer different perspectives (41, 42, 45). We therefore chose to understand how complex oscillation patterns arise by looking for experimental perturbations that could increase the occurrence of such events, instead of perturbations that change the distribution of oscillation frequencies.

Phosphoinositide metabolism plays an important role in recruiting regulators of Rho signaling cascade to plasma membrane (**Fig 2A**). We systematically knocked down major phosphatase involved in the phosphoinositide metabolism and scored for percentage of oscillations that exhibit complex oscillation patterns (**Fig 2B**). In wildtype (WT) cells, these complex patterns are rare, hence any conditions that leads to frequent appearance of complex pattern would be significant. We observed a significant increase in mixed-mode Rho oscillations in cells with knockdown (KD) of PTEN, a 3-phosphatase which catalyzes the dephosphorylation of the 3’ phosphate of PIP3 to yield PI(4,5)P2, but not with KD of Synaptojanin-2, which catalyzes the dephosphorylation of the 5’ phosphate of PI(4,5)P2 to yield PI(4)P (PTEN KD: n=17 experiments, 9/49 cells; Synaptojanin-2 KD: n=5 experiments, 0/15 cells) (**Fig 2C-D**). The most dramatic increase in complex oscillations occurs with knockdown of E-Syt1, which is a major membrane contact site protein in RBL cells and negative regulator of PI(4)P on the plasma membrane through lipid transfer from plasma membrane to ER (46-48). Majority of Rho oscillations in E-Syt1 knockdown displayed mixed-mode oscillation (18 experiments, 21/45 cells) (**Fig 2E**). Importantly, the mixed-mode oscillation in PTEN or E-Syt1 knockdown cells have much sharper nested pulses compared to wildtype cells, which can be captured in wavelet analysis as two separate peaks in the high and low frequency regimes (**Fig 2D-E, Supplemental Figure 1**).

**Fig 2.**
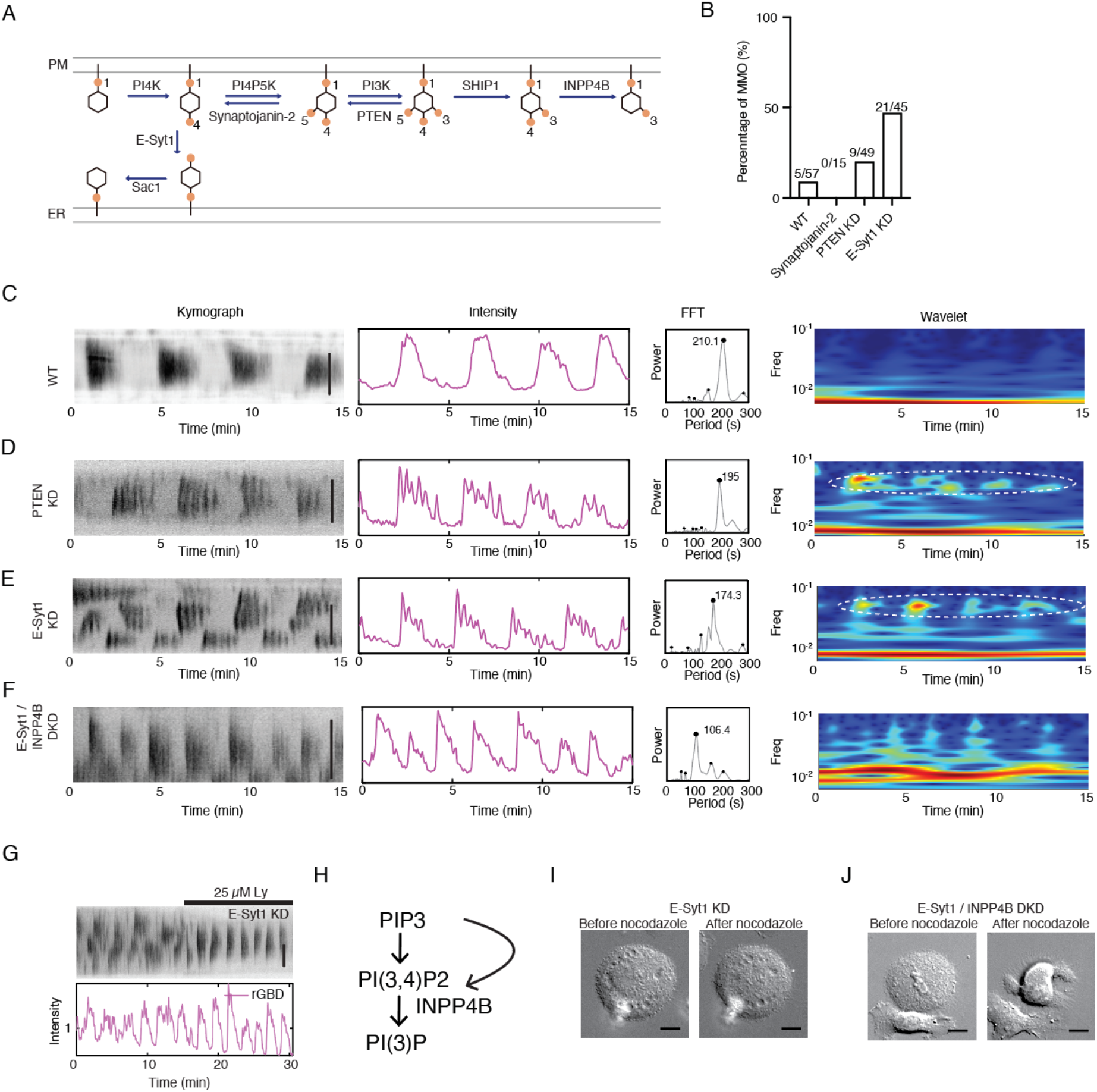
Complex mixed-mode Rho oscillations are upregulated in PTEN or E-Syt1 knockdown cells. (**A**) Schematic of the phosphoinositide metabolism pathways. (**B**) Bar plot showing percentage of mixed-mode Rho oscillations in wildtype (WT) cells (5/57 cells), Synaptojanin-2 KD (0/15 cells), PTEN KD (9/45 cells) and E-Syt1 KD (21/45 cells) conditions. (**C-F**) Representative kymographs, intensity profile, FFT and wavelet analyses of Rho oscillation in WT cells, PTEN KD cells, E-Syt1 KD and E-Syt1/INPP4B DKD cells. High frequency peaks in PTEN KD and E-Syt1 KD cell illustrated by wavelet analyses are circled with dotted line. (**G**) Representative kymographs and intensity profile of Rho oscillation in E-Syt1 KD cells treated with Ly294002. (**H**) Schematic representation of PIP3 metabolism to PI(3)P. (**I**) Representative DIC image of E-Syt1 KD cells before and after adding nocodazole. (**J**) Representative image of E-Syt1/INPP4B DKD cells before and after nocodazole. Horizontal and vertical scale bars: 10 μm.

Because PTEN is a negative regulator of PIP3, we hypothesize that the high frequency nested peaks are PIP3-dependent. PIP3 could regulate these high frequency oscillations either by regulating the synthesis or degradation of PI(3,4)P2. We next performed double knockdown (DKD) of E-Syt1 and SHIP1, the enzyme responsible for converting PIP3 to PI(3,4)P2, or INPP4B, the enzyme that degrades PI(3,4)P2. Mixed-mode oscillations were eliminated in E-Syt1/INPP4B DKD cells but not in E-Syt1/SHIP1 DKD cells. All the E-Syt1/INPP4B DKD cells imaged displayed simple patterns where high frequency peaks were reduced or disappeared in the wavelet analysis (4 experiments, 6/6 cells), indicating INPP4B is responsible for the sharp nested peaks **(Fig 2F)**. Similarly, lowering PIP3 had a smoothing effect on the high frequency nested peaks. When Ly294002, an inhibitor of PI3K, was applied to E-Syt1 KD cells displaying nested frequencies, Ly294002 caused disappearance of sharp peaks instantly (**Fig 2G**). These suggest that the high frequency peaks are regulated by PIP3/INPP4B pathway (**Fig 2H**), where high level of PIP3 leads to faster degradation of PI(3,4)P2 and sharper peaks, while low level of PIP3 leads to slower degradation of PI(3,4)P2 and merging of multiple high frequency peaks, which appears as single peaks with long duration. A clear phenotypic difference was also found between E-Syt1 KD cells and E-Syt1/INPP4B DKD cells. E-Syt1 KD cells were less contractile when microtubule was depolymerized by nocodazole. Their cell shapes and adhesion areas were much greater and did not change much before and after nocodazole treatment (**Fig 2I**). In comparison, E-Syt1/INPP4B DKD cells, similar to WT cells, contracted and reduced in footprint sizes significantly (**Fig 2J**). Together, these results suggest a strong correlation between Rho oscillation patterns with the outcome of cell contractility, where persistent Rho signals predict more productive contractility while short bursts with similar total durations were less effective. The most common form of mixed-mode oscillation consists of large-amplitude oscillations which are intercalated by small-amplitude oscillations, with three to four intercalated peaks were the most common (28/87 oscillations) (see examples in **Fig 2D and 2E**). A less common but reproducibly observed form of mixed-mode oscillation consists of large-amplitude oscillations intercalated by double peaks (3/87 oscillations) (**Fig 3C**). The duration between the major peak of one oscillatory cycle to the next is approximately 120 s while the peak-to-peak duration of the small-amplitude oscillations within the major peak’s oscillatory cycle is approximately 30 s (**Fig 3C and Fig 3D**). Again, most rise phases of either mixed-mode oscillations or double peaks are similar, all approximately 20 s. To determine whether a period-doubling bifurcation occurred (49), we also generated return map by plotting the interpeak interval (P1) relative to the next interpeak interval (P+1). Simple oscillations are defined as have only a single cluster on the Poincare plot (56/87 oscillations) (**Fig 3A and 3B**). Period-doubling bifurcation can be defined to have taken place if negative slope of the linear regression in a return map can be found (**Fig 3C-D**). Appearance of these period-doubling events (alternating between fast and slow cycles) indicating that a second stable oscillatory state likely exists.

**Fig 3.**
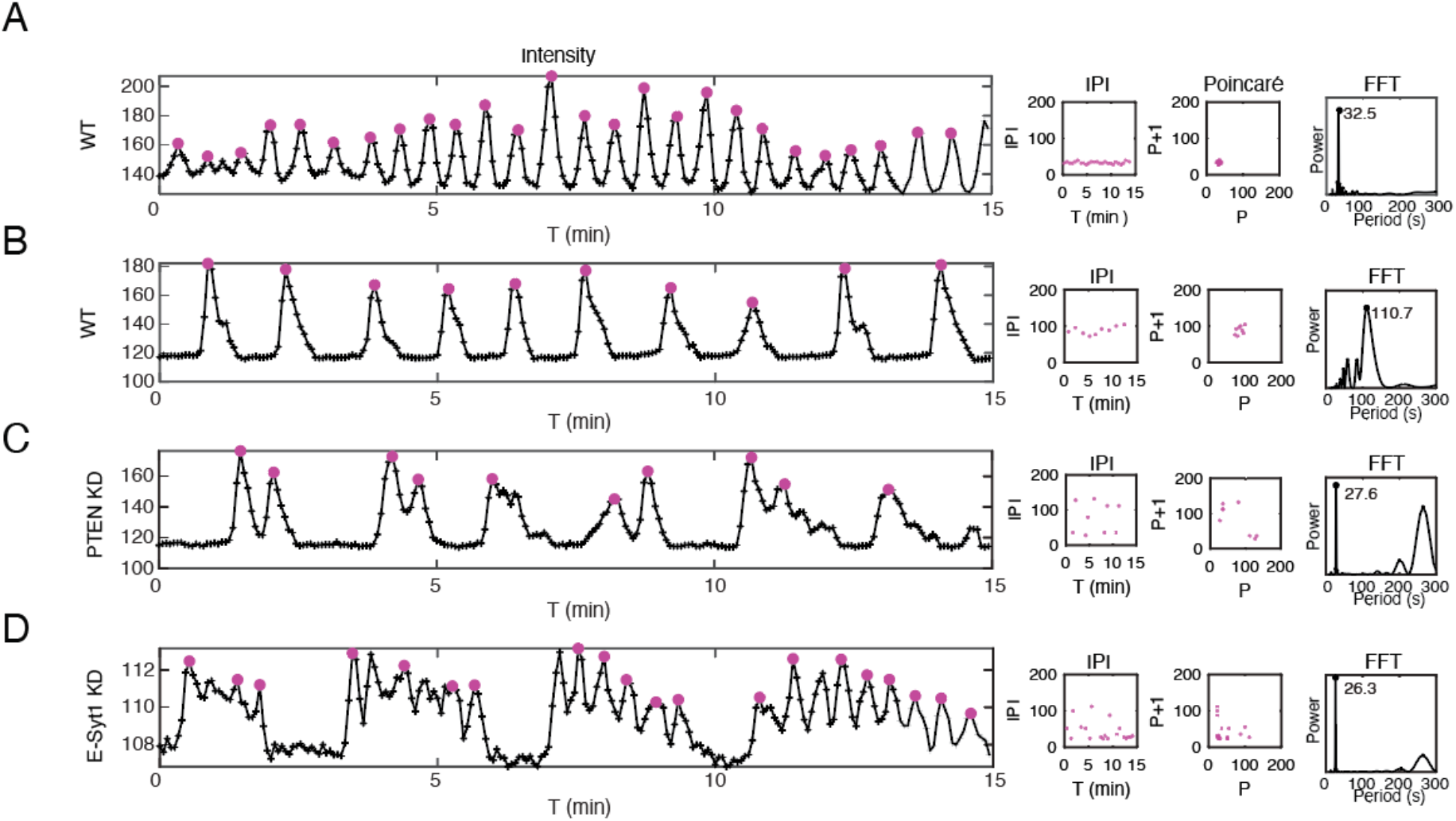
Line intensity profile, Interpeak interval (IPI), Poincare (P vs P+1) analysis and FFT of representative movies showing various simple and complex oscillations. Intensity profiles shows peaks selected for IPI and poincare analyses. FFT of all plots for both simple and complex Rho oscillations showed the major peak. (**A-B**) Simple oscillations with different periodicities (56/87 oscillations). (**C**) Mixed-mode oscillation of large-amplitude oscillations which are intercalated by double peaks (3/87 oscillations). (**D**) Mixed-mode oscillations which are intercalated by small-amplitude oscillations, with three to four intercalated peaks (28/87 oscillations).

To understand the slow oscillatory reaction, we first applied chemical inhibitors to determine whether the period of slow oscillations can be modulated (**Fig 4A**). We examined the effect of Ly294002 in both complex and simple oscillations. While Ly294002 consistently lead to smoothing of high frequency nested peaks in mixed-mode oscillation, effect of Ly294002 on the major period of oscillation was not consistent. Ly294002 slowed down major period in complex oscillations of PTEN KD cells (**Fig S2A**) while increased oscillation frequency for simple oscillations (7 experiments, 6/12 cells) **(Fig 4B**). We next perturbed PI(4)P metabolism by the pharmacological inhibition of PI4K using PI4Kα specific inhibitor GSK-A1. Inhibition of PI4K reduced the frequencies of simple oscillations (5 experiments, 9/9 cells). In simple oscillations with period of ∼60 sec, inhibiting PI4K leads to increased oscillation period to ∼150 sec (**Fig 4C**). This effect was seen in both WT cells, as well as PTEN KD cells that often display mixed-mode oscillations. Similar observations were seen in E-Syt1 KD cells. When GSK-A1 was added to cells displaying complex Rho oscillations, mixed-mode oscillation first transited into simple oscillations, followed by a reduction of Rho oscillation periodicity until the oscillations completely stopped (n=7 experiments, 8/8 cells) (**Fig S2B**).

**Fig 4.**
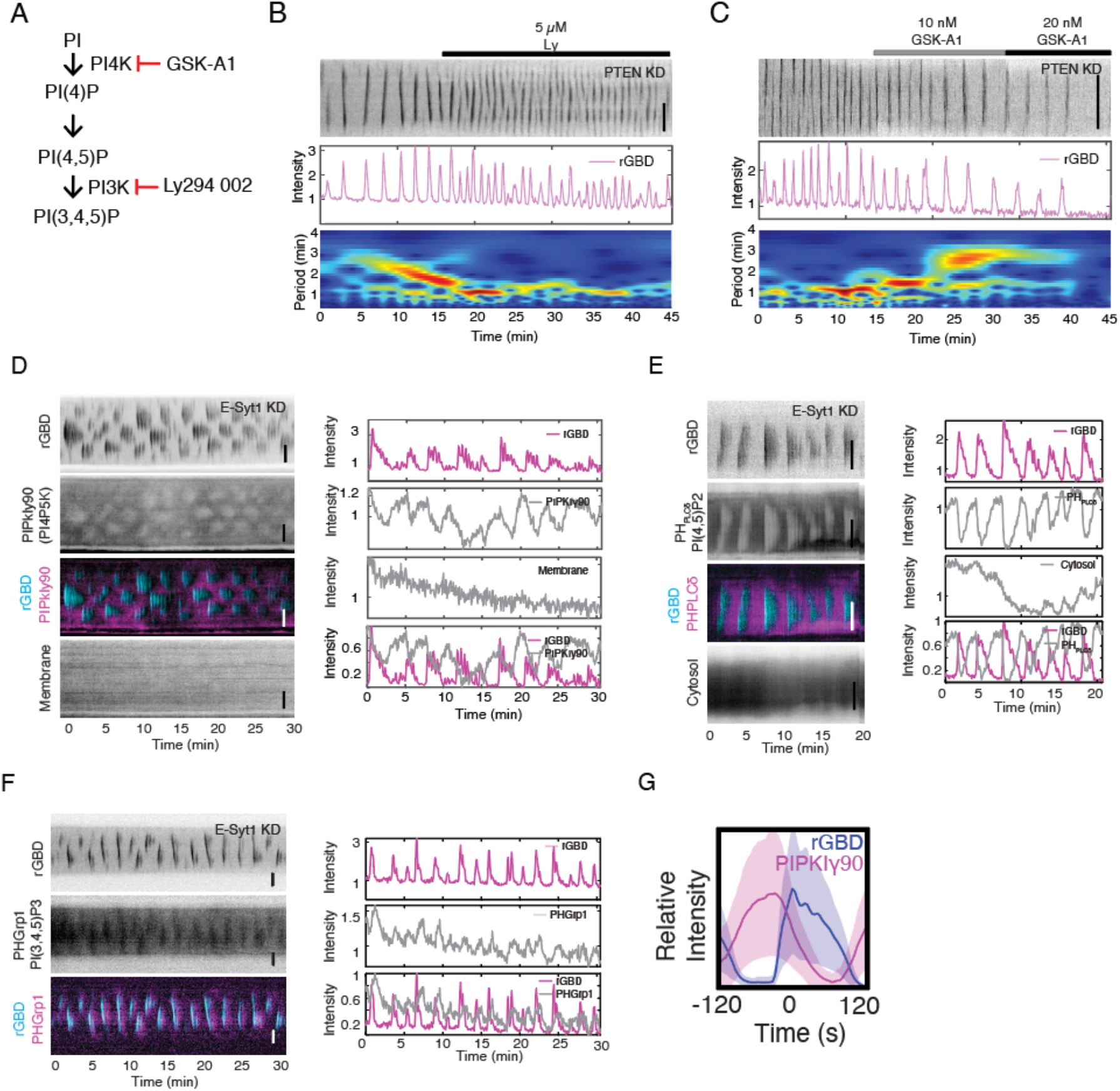
PI4P-PI4P5K pathway regulate the frequency of slow Rho oscillations. (**A**) Schematic of PI, PI(4)P and PIP3 metabolism. PI3K is inhibited by Ly294002 and PI4K is inhibited by GSK-A1. (**B**) Representative kymograph, intensity profile and wavelet analysis of Rho oscillations in a PTEN KD cell. Addition of PI3K inhibitor Ly294 002 increases Rho oscillation frequency (7 experiments, 6/12 cells). (**C**) Representative kymograph, intensity profile and wavelet analysis of Rho oscillations in a PTEN KD cell. Addition of PI4K inhibitor GSK-A1 reduces Rho oscillation frequency (5 experiments, 9/9 cells). (**D**) Representative kymograph and intensity profiles of Rho in E-Syt1 KD cell co-imaged with PIPKIy90 (PI4P5K) and membrane marker (3 experiments, n=3 cells). (**E**) Representative kymographs and intensity profiles of Rho in E-Syt1 KD cell co-imaged with PH_PLCδ_(PI(4,5)P2) and cytosol (2 experiments, n=2 cells). (**F**) Representative kymographs and intensity profiles of Rho in E-Syt1 KD cell co-imaged with PHGrp1 (PI(3,4,5)P2). (**G**) Average profiles by of aligned peaks of Rho and PIPKIγ90. The solid lines represent the mean intensities of the profile and shaded region represents the standard deviations of the intensities. Vertical scale bar: 10 μm.

The possibility that PI(4)P synthesis could regulate the periodicity of Rho oscillation is surprising because oscillation period is typically regulated by the lifetime of inhibitor in the activator-delayed inhibitor model. PI(4)P could directly recruit Rho or PI(4)P could serve as a precursor for PI(4,5)P2, which according to any prior knowledge, should be the activator. To better understand how PI4K regulates slow oscillation cycles, we imaged PI4P5K, which converts PI(4)P into PI(4,5)P2. Interestingly, PI4P5K as monitored by PIPKI γ90 levels were depleted when Rho was activated (**Fig 4D**). This depletion was not caused by the loss of TIRF signal due to membrane fluctuations as the same depletion was not seen by a CAAX membrane marker concurrently imaged in the same cell (3 experiments, n=3 cells) (**Fig 4D**). Consistent with the depletion of PI4P5K in sync with Rho activation, an anti-coupling was observed for PI(4,5)P2 and Rho, suggesting that the level of PI(4,5)P2 was also depleted when Rho activity was elevated (**Fig 4E**). Rho-induced contractility could induce membrane bending and loss of TIRF signals, but here the loss of TIRF signal resulting from membrane fluctuations was much less compared to the reduction of PI(4,5)P2, indicating the depletion was real (2 experiments, n=2 cells) (**Fig 4E**). In contrast, when we co-imaged PH_Grp1_, a sensor that binds to PIP3, we found that PIP3 was elevated relative to basal level (**Fig 4F**). In addition, nested frequency region correlated with higher PIP3 than regions with single pulses (**Fig 4F**). Collectively, this suggested a rather unexpected signaling cascade, where PI(4,5)P2 is depleted during Rho oscillation, and the replenishment of PI(4,5)P2 from PI(4)P regulates the oscillation period. Depletion of PI(4,5)P2 likely indicates that the consumption of PI(4,5)P2 (PI(4,5)P2->PIP3, PI(4,5)P2-> PI(4)P, etc.) exceeds its production (PI(4)P -> PI(4,5)P2, PIP3->PI(4,5)P2, etc.), which could be explained by the depletion of PI4P5K as Rho rises. While this may seem to rather counter-intuitive given the known positive association between PI4P5K and Rho, it does not contradict such relationship because there was still significant overlap between PI4P5K and Rho in time (**Fig 4G**).

## Conclusion

Paradoxical regulatory loops underlying oscillations have been increasingly recognized as important factors in the organization of signaling events. In this paper, we showed two such circuits underlying the spatiotemporal dynamics of lipid metabolism can govern Rho oscillations. The first major progress we made was to recognize that the heterogeneity of Rho dynamics cannot be explained by a single negative regulator. Mixing data with simple and complex oscillation initially lead us to wonder whether inhibiting PI3K leads to both increase and decrease of oscillation frequency, and this path turned out to be futile. Complex oscillatory phenomena, such as bursting or deterministic chaos that are neither periodic nor random, may arise through the coupling of two or more instability-generating mechanisms, each of which is capable of producing sustained oscillations on its own (45, 50). Although complex oscillations have been predicted in theory (42, 43, 51, 52) and extensively studied in chemistry and enzymatic reactions in the 1970s-90s, including Belousov–Zhabotinskii reaction (53, 54) and peroxidase-oxidase reaction (55, 56), there are little experimental studies in this direction in living cells (57-59). One possible reason is that unlike chemical or biochemical reaction conducted in the test tubes, where robust limit-cycle oscillations can be achieved, therefore deviation from perfect oscillations can be easily recognized, oscillations in living cell by default appear to be noisy. Thus, it is not always straightforward to differentiate between a noisy timeseries regulated by a simple feedback loop, and true chaotic dynamics governed by coupled networks. In addition, because the dynamic behavior is very sensitive to small parameter changes, genotypically identical cells can exist in different dynamical states, further complicating the single cell studies.

Upon recognizing the necessity to classify oscillations by simple and complex oscillations, the next challenge was to identify conditions that could bring about transitions from complex mixed-mode to simple oscillation and *vice versa*. We identified two such conditions. First, we show that PTEN knockdown increases complex mixed-mode behaviors of Rho oscillations. This is at least in part due to increased PIP3 levels because inhibiting PI3K in PTEN knockdown cells can reduce the complex oscillations. Second, we found that knockdown of E-Syt1, a protein anchored to the ER to mediate tethering to the plasma membrane and facilitate PI(4)P exchange from the plasma membrane to ER, can enhance mixed-mode oscillatory behaviors in more cells than PTEN knockdown cells. Intriguingly, while PI(4)P, PI(4,5)P2 and PIP3 are metabolically linked, knockdown SJ2, which should elevate PI(4,5)P2, did not have similar effects, highlighting the non-linear effect in the network, and the reason for this discrepancy would become apparent later.

Two commonly employed biochemical realization of oscillations are the activator-inhibitor scheme and the activator-depleted substrate scheme (60). In the activator-inhibitor model, oscillation occurs when the antagonist is too slow to immediately cancel out the activation; in the activator-depleted substrate model, oscillation occurs when the rate of substrate production could not maintain the rate of activator production. In either model, the refractory period follows the phase of activator during which time the inhibitor is degraded, or the substrate is replenished. The analysis of the elementary reactions involved here has not been completed, but our analysis is consistent with the activator-inhibitor model for the fast reaction, while the activator-depleted substrate model is involved in setting the slow reaction **(Fig 5A)**. In wildtype cells, the fast reaction is hard to resolve. Using these PTEN knockdown or E-Syt1 knockdown-induced mixed-mode oscillation, we found that the fast reaction oscillates at the time scale of around 30 sec, and correlate with high PIP3. In our previous study, we showed that PI3K plays a key role in governing cortical Cdc42 oscillations (∼ 30 sec) and inhibition of PI3K decreases Cdc42 oscillation frequency (32). The mechanism involved a delayed negative feedback (PIP3 -> SHIP1--| PIP3). Here a similar incoherent feedforward loop (PIP3-> INPP4--| PI(3,4)P2) is the most consistent with our results where PIP3 activates the turnover of PI(3,4)P2. For the slow reaction (oscillating in 1-2 min), we found that PI(4,5)P2 was depleted and reducing PI4K activity reduce oscillation period. Our results therefore are consistent with an activator-depleted substrate model where synthesis of PI(4,5)P2 from PI(4)P is slower than conversion of PI(4,5)P2 to PIP3, leading to its depletion, and the rate of PI(4)P replenishment regulates the reaction speed. When the fast and slow reaction are coupled, the slow reaction will essentially change the control parameter for the fast reaction, leading to either period doubling or mixed-mode oscillation. Rho was found to bind to several other phosphoinositides using a PIP membrane (28). Of which, PI(4)P binding to Rho appears to be the strongest. How the same PI(4)P->PI(4,5)P2->PIP3->PI(3,4)P2 signaling network support two circuits at different timescale remains to be established, which might involve different PI4K IIIα complexes (61). A parallel pathway involving PI(4)P-> PI(3,4)P2, which bypasses PIP3, could not be eliminated. However, because the high abundance of PIP3 is complex oscillation but PI(3,4)P2 is hardly detectable, we speculate there are at least two pathways that involves PIP3. Because two feedback loops of fast and slow time scale organized in tandem or in parallel can potentially generate chaos (62), the phosphoinositide network could offer more variations besides what was explored here.

**Fig 5.**
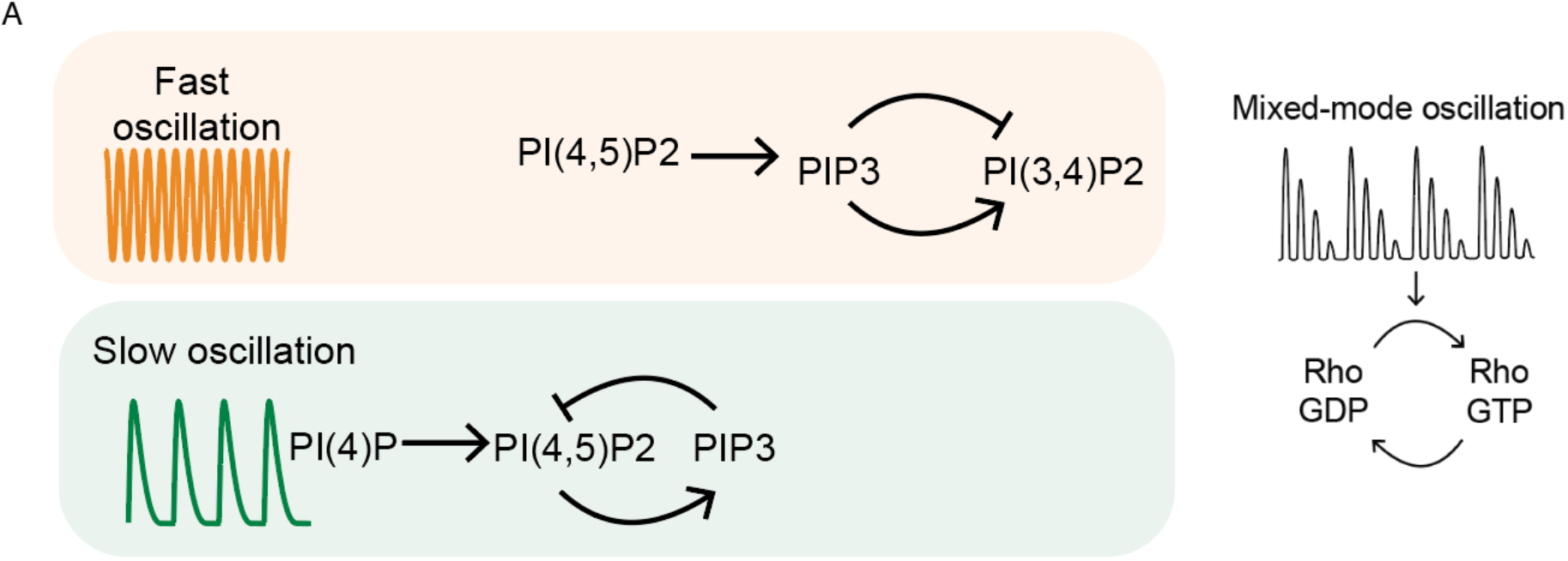
Topological representation of phosphoinositide metabolism that regulate fast and slow Rho oscillations.

Our main findings suggest an important role of lipid metabolism in regulating Rho dynamics. Because Rho oscillations was triggered by nocodazole that depolymerizes microtubule, our work also suggest that mitotic cortex is poised at the edge of chaos and microtubule could play a role in regulating the state transition from simple to complex oscillation or between any different dynamical states. Application of this framework is unlikely limited to mitotic cortex. Ultimately, one may understand the butterfly effect where immense phenotypic changes occurred due to a single mutation in PI3K (63).

## Methods

### Materials

The following reagents were purchased from commercial sources: Nocodazole, GSK-A1 and Ly294002 from Millipore Sigma. E-Syt1 shRNA in pRS vector (Cat # TR710045A, CCAAGTTCACTTGAGGTTAGAATGGCTAT and TR710045B, TTCCTTCGGACGCCGCTTGTTGGTGCTGG), PTEN shRNA in pRS vector (Cat # TR711414A CTTGACCAATGGCTAAGTGAAGACGACAA and TR711414B, ATAGAGCGTGCGGATAATGACAAGGAGTA), Synaptojanin 2 shRNA in pRFP-C-RS vector (cat. no. FI732825, TGTGCCTCTGCGGCAGCACCAGGTGAACT and FI732826, TTGTGGAGACAGAGCAGGCGATTTACATG) were purchased from Origene. Constructs of the following proteins were kind gifts: GFP-PTEN from Peter Devreotes (John Hopkins University School of Medicine), GFP-PIPKIγ90, mCherry-PH-Grp1 and iRFP-PHPLCδfrom Pietro De Camilli (Yale School of Medicine). mCherry-PIPKIγ90 was prepared by subcloning PIPKIγ90 into mCherry-C1 using restriction sites BgIII and EcoRI. GFP-rGBD was purchased from Addgene (Cat #26732).

### Cell culture and transfection

RBL-2H3 cell (ATCC) were cultured in MEM (Life Technologies) supplemented with 20% FBS (Sigma) and harvested with Trypsin-EDTA (0.25%) (Life Technologies). 1.2 × 10^6^ cells were transfected with 1 μg DNA or shRNA for each construct using the Neon transfection system (Life Technologies) following the manufacturer’s instructions (1200 pulse voltage, 20-ms pulse width and 2 pulse number), plated in two 35-mm glass bottom culture dishes (MatTek). For knockdown experiments, 1 μg/mL puromycin (Gibco) was added 24 hours post transfection for selection and imaged after 48 hours. Before imaging, media was replaced with imaging buffer Tyrodes [20 mM HEPES (pH = 7.4), 135 mM NaCl, 5.0 mM KCl, 1.8 mM CaCl_2_, 1.0 mM MgCl_2_ and 5.6 mM glucose]. To trigger rho oscillations during metaphase, 500 nM Nocodazole was added. For inhibitor experiments, drugs were diluted and added at final concentrations between 10 – 20 nM for GSK-A1 and 1 – 200 nM for Ly294002.

### TIRFM

TIRFM was performed using a Nikon TiE inverted microscope with 3 laser lines (488 nm, 561 nm, 642 nm). The microscope was equipped with an iLas2 motorized TIRF illuminator (Roper Scientific) and with a Prime95b sCMOS camera (Photometrics). All images were acquired using Nikon objectives (Apo TIRF 100×, N.A. 1.49 oil; 60×, N.A. 1.49 oil). Cells were imaged in two or three channels by sequential excitation with laser at 491 nm, 561 nm or 642 nm through a quad-bandpass filter (Di01-R405/488/561/635, Semrock). The microscope was controlled by Metamorph software (Universal Imaging). All images were acquired at 37 °C throughout the experiments using an on-stage incubator system (Live Cell Instrument, Seoul, South Korea).

### Image analysis

Post-acquisition image analysis was performed by Fiji and Matlab (Mathworks). A “reslice” tool and “average” projection filter in Fiji were used to generate kymographs. Auto-peak alignment, auto-correlation analysis, wavelet analysis and Fast Fourier Transformation were performed in Matlab. Custom codes are deposited on github (https://github.com/min-wu-lab/mmo-analysis). Raw data used for plotting in this paper has been deposited to Mendeley Data (doi: 10.17632/jzyp2w6hzt.1). To determine interpeak interval, individual peaks were identified based on secondary derivatives. Individual peaks were aligned at peak position and average curves were generated to obtain rise and drop phases. To determine whether a period-doubling bifurcation occurred, the return map was generated by plotting the interpeak interval (P1) relative to the next interpeak interval (P+1).

## Supplemental Figure

**Fig S1.**
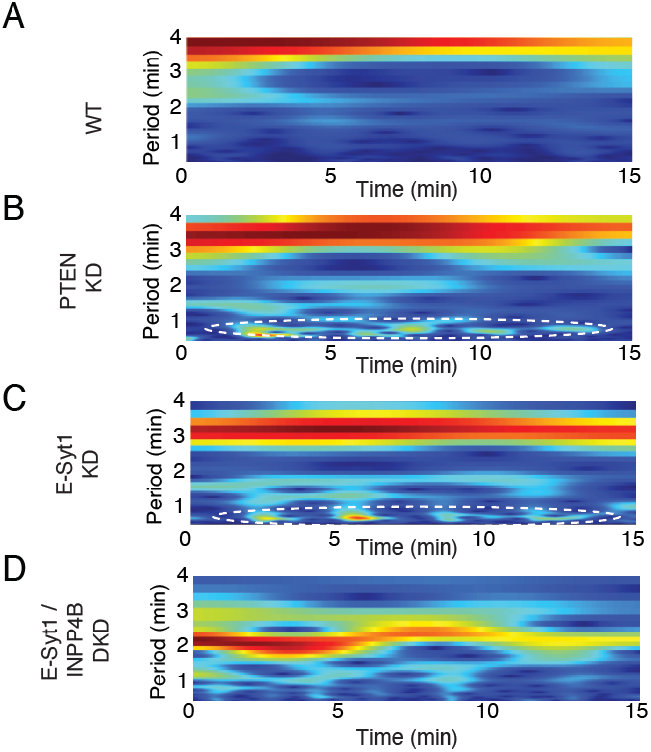
Complex mixed-mode Rho oscillations are upregulated in PTEN or E-Syt1 knockdown cells. Wavelet analyses of Rho oscillation in WT cells, PTEN KD cells, E-Syt1 KD and E-Syt1/INPP4B DKD cells plotted as time vs period. The wavelets correspond to wavelet plotted as time vs frequency in **Fig 2C-F**. High frequency peaks in PTEN KD and E-Syt1 KD cell illustrated by wavelet analyses are circled with dotted line.

**Fig S2.**
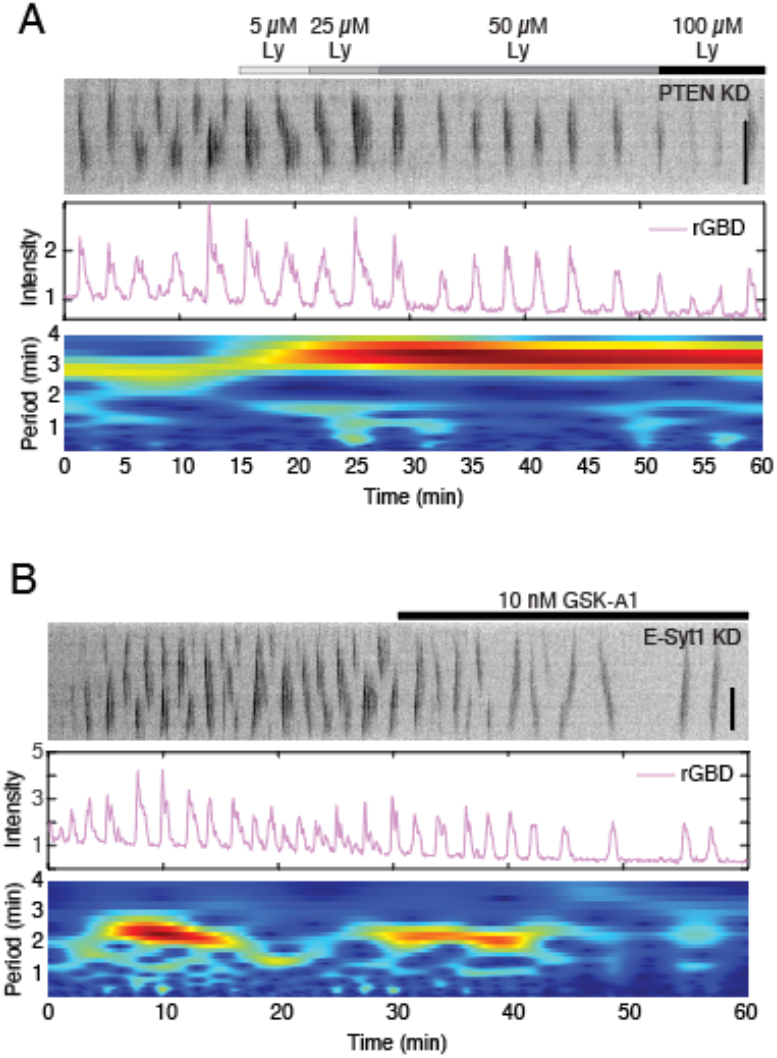
Inhibition of PI3K or PI4K converts mixed-mode oscillation to simple oscillation and affect the major cycle time. (A) Representative kymograph, intensity profile and wavelet analysis of Rho oscillations in a PTEN KD cell before and after addition of PI3K inhibitor Ly294 002. (B) Representative kymograph, intensity profile and wavelet analysis of Rho oscillations in a E-Syt1 KD cell before and after addition of GSK-A1.

